# Designing oil palm landscapes to retain biodiversity using insights from a key ecological indicator group

**DOI:** 10.1101/204347

**Authors:** Claudia L. Gray, Eleanor M. Slade, Darren J. Mann, Owen T. Lewis

**Author notes:** Current address: EDGE of Existence, Conservation Programmes, Zoological Society of London, Regent’s Park, London, NW1 4RY. Corresponding author: Claudia Gray.

## Abstract

Oil palm expansion threatens biodiverse ecosystems across the tropics. However, palm oil is a widely used and profitable crop, so identifying strategies that mitigate the impact of oil palm expansion on biodiversity is important. Riparian reserves (strips of forest along rivers) are protected in many countries for hydrological reasons and also support species that would not otherwise persist in oil palm. However, management guidelines for riparian zones have been informed by relatively few ecological studies. We assessed how the structural features and landscape context of riparian reserves in Sabah, Malaysia affected dung beetle communities. We also tested the use of flight intercept traps to study movement of dung beetles along linear forest corridors. Overall, dung beetle abundance in riparian reserves was 54% lower than in logged forest areas, but all species observed in the logged forest were found in at least one riparian reserve site and both species richness and diversity increased with reserve width. Distance from a large block of continuous forest affected dung beetle community composition but not species richness, abundance, or functional diversity. The amount of forest cover in the surrounding landscape improved the retention of species within riparian reserves, and increases in vegetation complexity corresponded with higher functional richness and functional dispersion. The flight intercept traps did not indicate that there is net movement of individuals out of logged forest areas into the riparian reserves. The species richness of 30 m reserves (the suggested requirement of reserves in Sabah) was only 10% lower than in logged forest, but our data indicate that riparian reserves of at least 50 – 80 m are needed for species richness and diversity to equal that in nearby logged forest. These findings, particularly if they apply more widely to forest-dependent taxa, should be taken into account when setting policy and sustainability guidelines for oil palm plantations, both in areas undergoing conversion from forest and in existing oil palm plantations where forest restoration is required.

## INTRODUCTION

Tropical landscapes hold the majority of the planet’s biodiversity and are a priority for conservation action (Gardner *et al.* 2009). Agricultural expansion and intensification are amongst the major threats to biodiversity in tropical regions (Laurance, Sayer & Cassman 2014). Recent evidence suggests that since many tropical species are highly sensitive to increases in land use intensity, a policy of land sparing (intensifying land use to avoid conversion of undisturbed habitat) will be more successful than land-sharing (limiting intensification to maximise biodiversity retained within productive landscapes) (Edwards *et al.* 2014; Gilroy *et al.* 2014a; Wehrden *et al.* 2014). However, even if we successfully target intensification to spare land for nature, there will remain some habitat fragments within agricultural landscapes that can benefit biodiversity. Designing these fragments to maximise biodiversity conservation will remain important.

A widespread example of fragmented habitat retained even within intensive agriculture is riparian zone vegetation. Riparian reserves, or buffer zones (strips of non-crop habitat protected alongside rivers, streams and lakes) are protected in many countries because they reduce the flow of chemicals and sediment into the water, thereby limiting soil erosion, lowering flood risk and preserving water quality (REFS). A wide range of countries have legislation protecting riparian zone vegetation (McDermott, Cashore & Kanowski 2010). Even in a land-sparing scenario, it is recognised that these landscape features should be retained for the other ecosystem services they provide (Edwards *et al.* 2014).

In addition to their hydrological benefits, riparian reserves can support many terrestrial species that would not otherwise survive in areas of agriculture or plantation (Marczak *et al.* 2010; Gray *et al.* 2014, 2015). However, relatively little ecological information is available on how riparian zone management impacts terrestrial species, particularly in tropical regions. In addition, only a small number of the existing studies give clear management recommendations, and the methods used to make the recommendations are highly variable, so results can be contradictory. Improving the information available on terrestrial species is important for maximising the conservation value of riparian reserves, as guidelines based on aquatic species or hydrological processes are unlikely to be adequate for terrestrial fauna in riparian zones (Lee, Smyth & Boutin 2004; Viegas *et al.* 2014).

Oil palm is widely cultivated across SouthEast Asia and is expanding rapidly across the tropics (Butler & Laurance 2010; Wich *et al.* 2014; Gilroy *et al.* 2014b). As many species found in nearby forests are lost after conversion to oil palm (Savilaakso *et al.* 2014) the growth of the industry is a concern for conservationists. Global demand for palm oil is still increasing (FAOSTAT 2013), so identifying strategies to limit the negative environmental impact of oil palm plantations is of great importance. Setting appropriate guidelines for riparian reserves in these landscapes could help retain biodiversity that would otherwise be lost.

The number of meters of vegetation required on each side of the river is one of the clearest, most common specifications made for management of riparian areas. Whilst guidelines do also cover other features of the reserves (e.g. vegetation structure, chemical application), they are less commonly specified in policy (McDermott, Cashore & Kanowski 2010)(McDermott, Cashore & Kanowski 2010). A summary of riparian zone legislation for many key oil palm producing nations is given in Appendix 3 of (Barclay *et al.* 2017). In Indonesia, forest must be 50m wide on each riverbank for all rivers under 30m, and 100m on each side for all rivers over 100m. In Sabah, Malaysia, the legal requirement is that riparian vegetation is 20 m wide on each bank of any river more than 3 m in width (*Sabah water resources enactment* 1998). Elsewhere in Malaysia the requirements are between 5 and 40m depending on the size of the river. Riparian buffer zones in Ghana, one of the major oil palm producers in Africa, are between 10 and 60 m, again depending on the size of the river. The Roundtable on Sustainable Oil Palm (RSPO) requires that in the absence of any specific national guidelines, riparian reserves should be between 5 and 50 m for rivers up to 50 m wide (the required width of a buffer zone increases in 10m steps corresponding to each 10 m increase in width of a river) and then at least 100 m of buffer are recommended for rivers over 50 m wide (Barclay *et al.* 2017).

Dung beetles are a key ecological indicator group, as they are sensitive to habitat disturbance (Slade, Mann & Lewis 2011), their responses are congruent to those of other animal groups (Gardner *et al.* 2008) and they provide important ecological functions (Nichols *et al.* 2008). In addition, species specific trait data is available for many species, so that a range of functional metrics can be calculated in addition to traditional biodiversity metrics of species richness and alpha and beta diversity. Functional traits provide an insight into how species’ responses to land use change may be dependent on their life-history and ecological niches, and can reveal shifts in community composition that are not necessarily detected by traditional metrics (Mouillot *et al.* 2013). Recent studies in Borneo have shown that the response to conversion to oil palm in both vertebrate and invertebrate species differs with functional traits such as body size and tropic level (Senior *et al.* 2013), and that the functional diversity of dung beetles in particular is lower in oil palm than in logged forest (Edwards *et al.* 2013; Gray *et al.* 2014).

Here, we quantify the biodiversity impact of the landscape context and structure of riparian reserves, to inform policy for riparian zone management. We test the hypotheses that riparian reserves of greater width and vegetation complexity increase the extent to which riparian reserve fauna resemble communities in logged forest. We also assess how the amount of forest remaining in the area around the reserves, and the distance to the nearest point at which the reserve joins a large (>2000 ha) area of logged forest affects dung beetle communities. We examine whether the role of riparian corridors in facilitating dispersal through oil palm landscapes can be assessed using flight intercept traps. In conclusion, we present policy recommendations for riparian reserve width in oil palm landscapes in Sabah.

## METHODS

### STUDY SITES

All study sites were located within a 600 km^2^ area around and including the Stability of Altered Forest Ecosystems (SAFE) project site in Sabah, Malaysian Borneo (117.50 N, 4.60 E). The area is a mixture of twice-logged lowland dipterocarp rainforest, acacia and oil palm plantations, in which palms were planted between 2006 and 2012. Further details of the project area are given in Ewers et al. (2011) and further details on the invertebrate communities and vegetation characteristics of the riparian reserves we sampled are presented in (Gray *et al.* 2014, 2015; Gray & Lewis 2014; Luke *et al.* 2017b; a). All data collection was carried out between September and November 2012.

### EFFECTS OF RIPARIAN RESERVE STRUCTURE AND LANDSCAPE CONTEXT ON DUNG BEETLES

Dung-baited pitfall traps were set at 23 riparian reserve sites in oil palm and five riparian logged forest reference sites (Fig S1). For riparian reserves, we maximised the number of sites, range of reserve widths, and range of distances from the forest, whilst also ensuring that all sites were surrounded by oil palm plantations on both sides and at least 1 km apart; this spatial distribution allows the dung beetle assemblages at each site to be treated as independent samples (Larsen & Forsyth 2005).

Forest river sites were selected to achieve spatial interspersion with riparian reserve sites as far as possible. A number of studies making recommendations for riparian zone management have surveyed riparian zones within continuous areas of forest to set a reference level to which riparian zones in agricultural areas can be compared (e.g. (de Lima & Gascon 1999; Hagar 1999; Pearson & Manuwal 2001). We used logged forest comparison sites to set this reference as there was no primary forest within 50 km of our field sites. Riparian reserve forests in oil palm are usually remnants of degraded logged forests, and so it is unrealistic to expect that that the degraded riparian reserve forest should be managed with the goal of resembling undisturbed primary forest(Gray *et al.* 2014).

At each site we set five traps, each baited with 25g human dung and collected after 48 hours, following standard methods for surveying dung beetle communities (see (Gray *et al.* 2014) for further details). One trap was placed at each corner of a 50 m × 25 m rectangle (two traps approximately 1 m above the high water line and 50 m part, two traps 25 m from the high water line and 50 m apart), with one trap in the centre of the rectangle. This design ensured that all 5 traps were sampling the same community at each site (Larsen & Forsyth 2005) and were always within the riparian reserve vegetation. Traps were set at a maximum of two sites during each 48 hour period.

Dung beetle (Scarabaeidae, Scarabaeinae) specimens were stored in 90% alcohol and later identified using Balthasar (1963), Boucomont (1914) and the works on Bornean Scarabaeinae by Ochi and Kon (e.g. Ochi et al., 1996), the reference collections housed in the Oxford University Museum of Natural History (OUMNH) and the Natural History Museum, London. We combined data from all five traps at each site to calculate the following community metrics for a) the total complement of dung beetles and b) the subset of species endemic to Borneo (highlighted in Table S1): dung beetle abundance, biomass, rarefied species richness, alpha diversity (Shannon index), and beta diversity (species turnover between traps within a site; mean Sørensen’s similarity index). To calculate biomass, we weighed beetles from 24 species taken from across the whole range of body sizes (between 7 – 51 individuals per species, mean = 27, s.d. = 8) and used a polynomial regression to estimate biomass from body length measurements for the remaining species (Log_10_ (biomass) = −1.64 + 5.61*Log_10_(length) − 4.39*Log_10_(length)^2^ + 1.99*Log_10_(length)^3^, R^2^ = 0.982).

For the total complement of dung beetles in the pitfall traps at each site we also calculate three indices of functional diversity (Villéger, Mason & Mouillot 2008; Laliberté & Legendre 2010): a) functional richness (FRic), the total volume of the centroid in trait-space that is occupied by the species at each sampling point, b) functional dispersion (FDis), the average distance of species from the centroid, weighted by their relative abundances, and c) functional evenness (FEve), the evenness of the distribution and relative abundances of the species.

For each site we calculated a) the average width of the reserve, from measurements at both ends and the centre of the trap rectangle (widths are given for one side of the river, to match current policy terminology), b) distance to the point where the riparian reserve joined the nearest large (> 2000 ha) block of logged forest (both along the corridor and linear distance) and c) the proportion of forest cover within a buffer zone of radius 1 km around each trapping site. All these variables were calculated in a GIS (ArcMap version 10.1) using GPS points of the trap locations, tracks of the riparian reserve boundaries and a land cover map derived from a maximum likelihood supervised classification of SPOT satellite images combined with digitised maps of the plantations. Distances to the nearest large block of forest measured in a straight line versus along the corridor were very highly correlated (r = 0.90, df = 21, p < 0.0001), so for all analyses we used only distance along the corridor as this is more ecologically relevant for species that use the riparian forest. The area of forest in the 1 km radius buffer and width of riparian reserves were also correlated (r = 0.69, df = 21, p = 0.0002). Therefore, to measure how the forest in the 1km buffer zone varied from what we would expect based only on the measurement of width at the point of trapping, we regressed the square-root of the area of forest in the buffer against the width of the reserve and used the residuals for subsequent analyses.

To calculate a measure of vegetation complexity at each site, we measured humus depth, canopy density (using a spherical densiometer) and basal area (using the angle point method (Bitterlich 1984)). We estimated the height of the tallest tree to the nearest 5m using a ruler held at arm’s length and a known reference height at the base of the tree, and scored the under-storey vegetation density (below 2m) and mid-storey vegetation density (between 2m and 5m) on an ordinal scale of sparse (fewer than 20 stems or branches) medium (20 – 60 stems or branches) and dense (few patches of light and 60 – 100+ stems or branches). To obtain one numerical index summarising the greatest variation in the vegetation data, we carried out a metric scaling analysis on all the vegetation and soil measurements. The first axis of the analysis was positively correlated with canopy density, tree height, humus depth, basal area and mid-storey density and explained 54% of the total variation in the data. Since this output is therefore capturing variation in the 3-dimensional structure of the vegetation, we refer to it as a vegetation complexity index.

Having obtained all these variables, we ran linear models to test for a relationship between each of the dung beetle community metrics listed above and riparian reserve width, distance to the nearest logged forest, area of forest within a 1 km radius (relative to what would be expected for reserve width), vegetation complexity and all two way interactions apart from the interaction between width and distance to nearest logged forest (we did not have sufficient data to test for this interaction as we had no sites where the riparian reserve was wide and also far from the forest). Riparian reserve width was log transformed to meet model assumptions. Variables were removed from the model in order of least significance until the minimal adequate model was obtained.

For all cases in which there was a significant relationship between a particular dung beetle community metric and riparian reserve width, we used the model-predicted values to explore the sensitivity of the dung beetle community to changes in reserve width that could be implemented in policy. We took the mean and lower bound of the 95% confidence interval (CI) for each metric in logged forest sites and found the width at which species richness in riparian reserves matched the forest values. For all response variables where there was no effect of riparian reserve width, distance to the nearest logged forest, proportion of forest in the surrounding area or vegetation complexity, we carried out a subsequent analysis to test whether the communities across all riparian reserve sites differed from the logged forest reference sites.

To examine changes in community composition, we used a PERMANOVA (permutational multivariate analysis of variance with 1000 permutations). We tested whether riparian reserve width, distance to the nearest logged forest, proportion of forest in the surrounding area, and vegetation complexity explained differences in riparian reserve community composition of a) the total dung beetle dataset and b) the subset of Borneo endemics.

### RIPARIAN RESERVES AS CORRIDORS

We investigated beetle movement from the continuous logged forest into and along the corridors using flight intercept traps (FITs) at four spatially independent locations (all > 6 km apart) where a riparian reserve corridor joined the large area of logged forest. At each of these corridor junctions, FITs were set at 0 m, 200 m, 500 m, and 1 km from the point where the riparian reserve met the logged forest (Hill 1995)). For each trapping location we calculated riparian reserve width, proportion of forest cover in a 1 km radius and vegetation complexity (see above).

Each FIT was made from black nylon mesh (mesh size = 0.5 cm, dimensions = 1.5 m × 2 m), with ten collection trays (30 cm long, 20 cm wide, 10 cm depth) placed on the ground on each side of the net and filled with a solution of water, detergent and salt. All FITs were protected by a rain cover. Tray contents were collected separately for each side of the FIT every 48 hours for 6 days.

FITs were aligned perpendicular to the river, so that the trays on one side collected the insects intercepted by the net as they flew in a direction away from the forest, and the other side intercepted insects flying back towards the forest. Wherever possible, traps were placed across ‘natural paths’ in the forest, i.e. existing clearer sections through which insects were likely to be flying. No vegetation was cut or cleared.

For each FIT we calculated the abundance, biomass, species richness and alpha diversity (Shannon index) of dung beetles. We tested whether each of these was predicted by distance along the corridor, side of the trap (facing towards versus away from the forest) and the distance by side interaction using a linear mixed effects model with FIT location specified as a random factor.

All analyses were carried out in the R statistical software (version 3.0.2 (R Core Team 2014)) using the packages lme4 (Bates *et al.* 2014 p. 4) andvegan (Oksanen *et al.* 2013) and FD (Laliberte & shipley 2011).

## RESULTS

From the pitfall traps we identified 5775 individual beetles of 59 species (including 27 Borneo endemics, Table S1). From the FITs we identified 3306 individuals of 68 species (including 33 Borneo endemics, Table S2).

### EFFECTS OF RIPARIAN RESERVE STRUCTURE AND LANDSCAPE CONTEXT ON DUNG BEETLES

All dung beetle species found in the logged forest were recorded in at least one riparian reserve site. There was no significant effect of riparian reserve width, distance from forest, forest cover in a 1 km radius, buffer habitat complexity or any two way interactions on either the abundance or biomass of dung beetles (both overall and Borneo endemics subset, p > 0.06 in all cases). There was also no significant difference between dung beetle biomass in riparian reserves and logged forest for all species combined (F_1,26_ = 0.93, p = 0.34) or Borneo endemics (F_1,26_ = 3.58, p = 0.07), although the trend in the data for the endemics was that biomass in the riparian reserves was lower. However, dung beetle abundance in the riparian reserves was 54% lower than logged forest for all species (F_1,26_ = 10.48, p = 0.0033), and 50% lower for Borneo endemics (F_1,26_ = 13.35, p = 0.001, Table S3).

There was a significant positive effect of riparian reserve width on species richness for all species (F_1,21_ = 9.01, p = 0.007, Fig 1 a)), and for the Borneo endemics (F_1,21_ = 5.49, p = 0.029, Fig 1 b)). There was also a significant effect of both width and proportion forest cover in the surrounding landscape on the diversity of dung beetles overall (F_2,20_ = 8.64, p = 0.002, Fig 1 c) and Fig S3), but no significant effect of any explanatory variables on the diversity of endemic beetles (p > 0.11 in all cases). There was also no significant difference between the diversity of endemic species in logged forest and riparian reserves (F_1,26_ = 1.25, p = 0.27). Table 1 gives the widths at which the mean and lower 95% CI for species richness and diversity in the riparian reserves match the mean and lower 95% CI for the logged forest sites (corresponding to arrows in Fig 1). Table 2 shows the gain in species richness obtained by increasing the legally required 20m reserve width to 30, 50 and 80 m on each side of the river.

**Fig 1.**
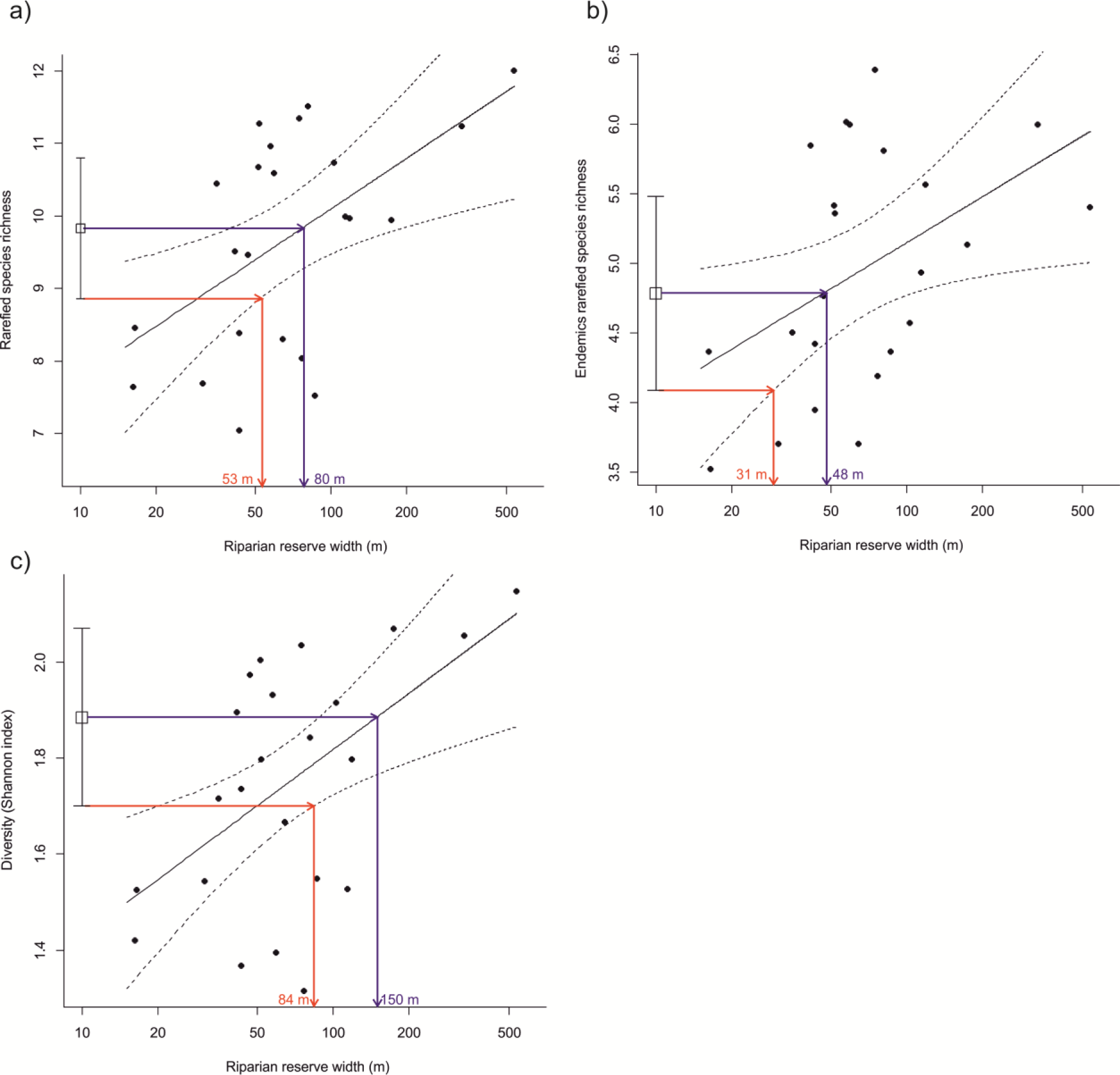
The species richness of all dung beetles (a), the species endemic to Borneo (b) and the diversity of all dung beetles (c) increase with riparian reserve width. Full circles show values for the riparian reserve sites. The solid and dotted lines show the model predicted values and 95% confidence intervals; unlogged width values are shown for ease of interpretation although the analysis was conducted on logged riparian reserve widths. Square and bars give the mean and 95% CI for the logged forest sites. As there was also a significant relationship between diversity and area of forest in the surrounding area, model predications in panel c) were obtained by specifying the average value for forest in the surrounding area.

**Table 1.** Riparian reserve widths at which mean and lower 95% CI for dung beetle community metrics is equal to corresponding values in logged forest

|  | Width* where<br>riparian reserve mean<br>= logged forest mean | Width* where<br>riparian reserve lower 95% CI<br>= logged forest lower 95% CI |
| --- | --- | --- |
| All species richness | 80 | 53 |
| Endemic species richness | 48 | 31 |
| All species diversity | 150 | 84 |
| Endemic species diversity | No relationship with width | No relationship with width |

\*width is given for one side of the river only, in metres, to match existing policy specifications

**Table 2.** Estimates of the biodiversity consequences of expanding riparian reserves*.

|  | 30 m<br>width | 50m<br>width | 80m<br>width | 150m<br>width | Logged<br>forest |
| --- | --- | --- | --- | --- | --- |
| upper 95% CI<br>species richness | 9.66 | 9.98 | 10.43 | 11.28 | 10.79 |
| species richness | 8.85 | 9.38 | 9.86 | 10.50 | 9.83 |
| lower 95% CI<br>species richness | 8.05 | 8.78 | 9.29 | 9.71 | 8.86 |
| % gain in species<br>compared to<br>30m buffer<br>(upper 95% CI) | 0% | 3.3% | 8.0% | 16.9% |  |
| % gain in species<br>compared to<br>30m buffer | 0% | 6.0% | 11.4% | 18.6% |  |
| % gain in species<br>compared to<br>30m buffer<br>(lower 95% CI) | 0% | 9.1% | 15.4% | 20.6% |  |

\*figures are derived from the model given in Fig 1 a) so the values given here also correspond to species richness rarefied to 33 individuals (the minimum number of beetles that occurred at any site).

There was no significant effect of any explanatory variables on total or endemic beta diversity (p > 0.15 in all cases), and no significant difference in beta diversity between forest and riparian reserves for all species (F_1,26_ = 1.28, p = 0.27) or the Borneo endemics (F_1,26_ = 0.89, p = 0.35).

There was a significant positive relationship between the vegetation complexity of the reserves and both functional richness (FRic, F_1,21_ = 5.0, p = 0.036, Fig 2 a)) and functional dispersion (FDis, F_1,21_ = 7.85, p = 0.011, Fig 2 b)). However, there was no relationship between any explanatory variables and functional evenness (FEve) (p > 0.06 in all cases). FEve also did not differ between logged forest and riparian reserves (F_1,26_ = 3.0779, p = 0.09).

**Fig 2.**
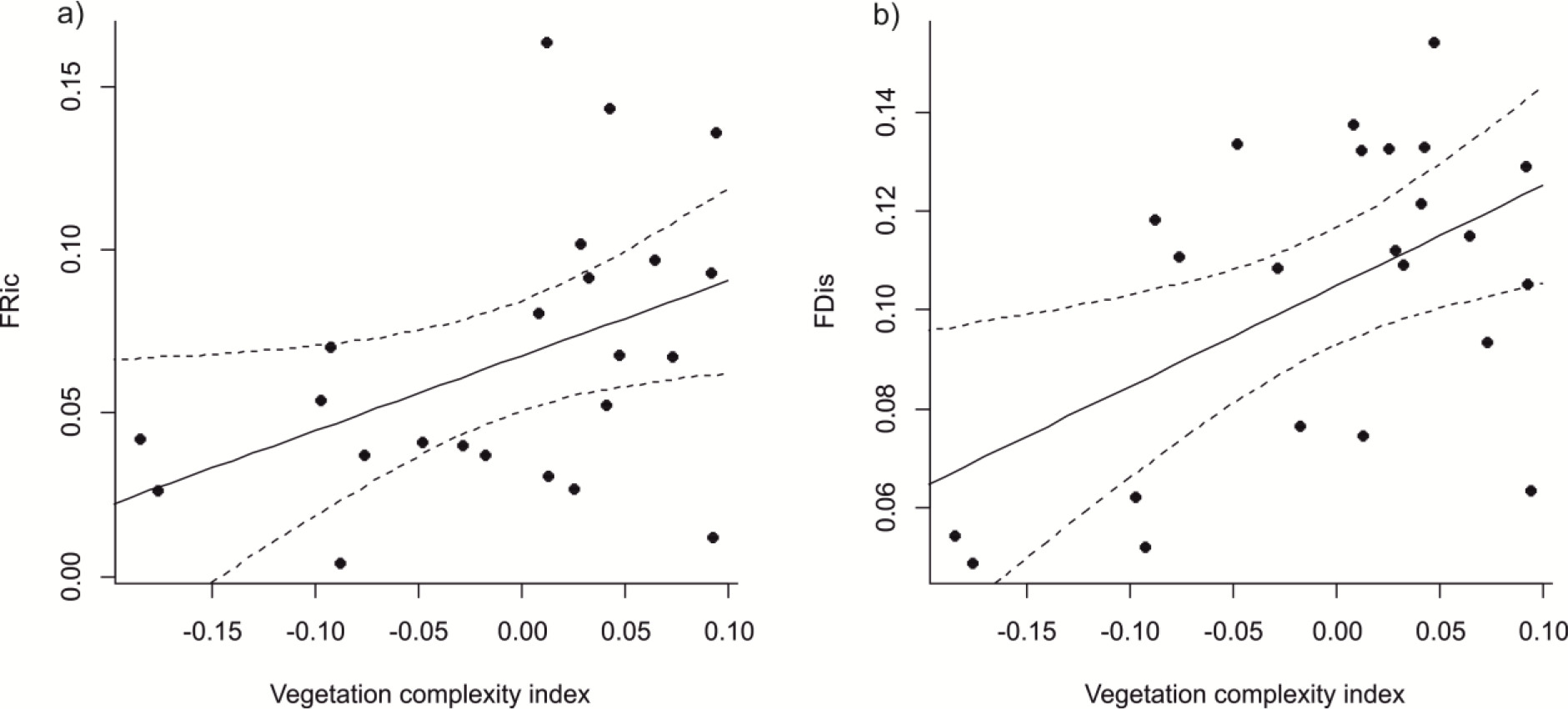
Functional richness, FRic (a) and functional dispersion, FDis (b), increase with the vegetation complexity within riparian reserves.

There was a significant change in community composition with distance from the nearest logged forest (F_1,21_ = 2.65, p = 0.03, Fig. S2) for all dung beetle species combined, but no relationship between the composition of the endemics and any explanatory variables (p > 0.1 in all cases).

### RIPARIAN RESERVES AS CORRIDORS

There was no significant effect of distance to the nearest logged forest, trap side or their interaction on any of the dung beetle community metrics calculated from the FITs (Fig 3).

**Fig 3.**
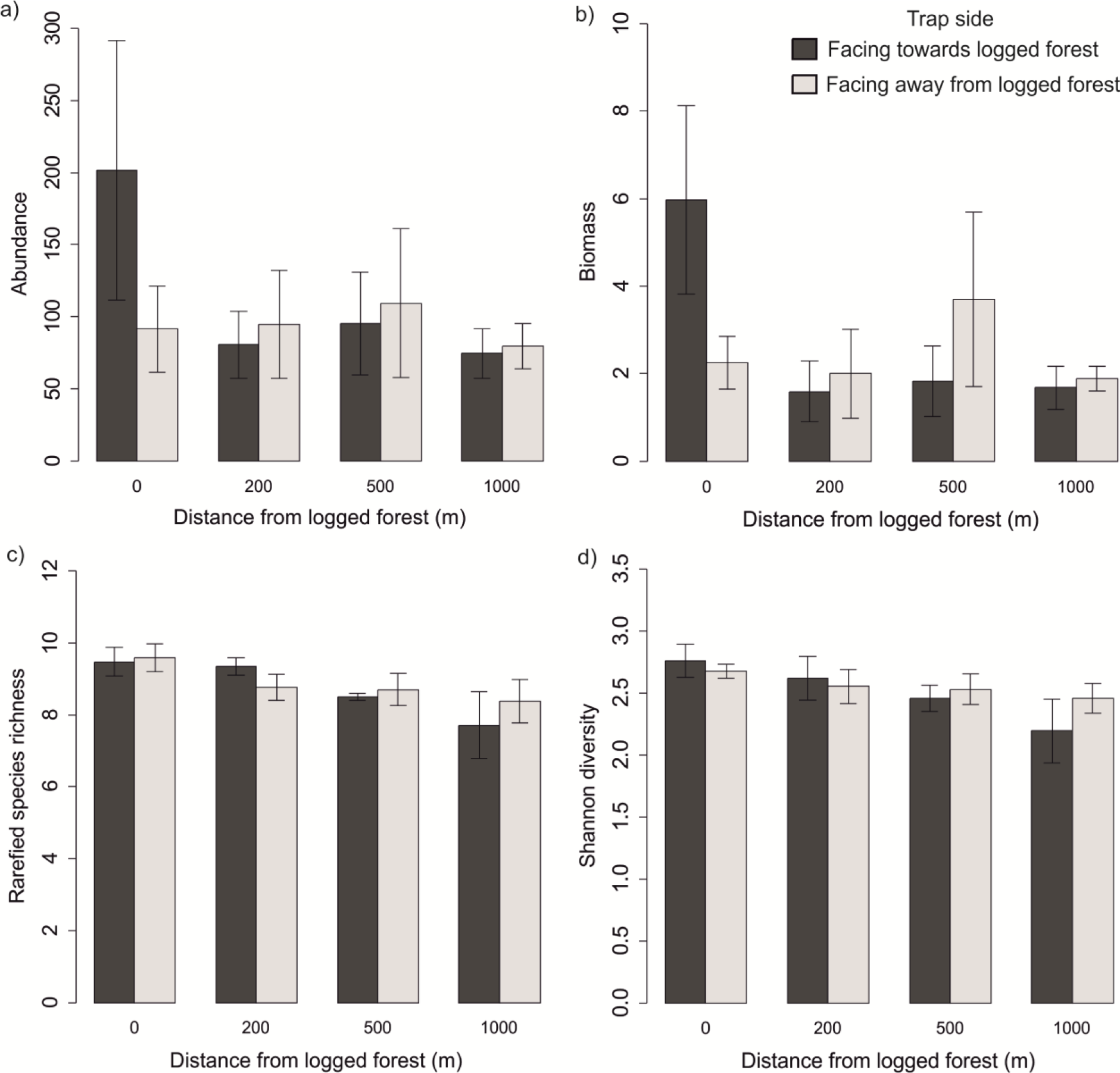
Abundance (a), biomass (b), species richness (c) and diversity (d) of dung beetles (± s.e). caught in FITs at increasing distances from logged forest along the riparian corridor.

## DISCUSSION

As oil palm plantations expand across the tropics, maximising the biodiversity retained within these landscapes will be important. Our results suggest that increasing the width of riparian reserves and minimising distance to remaining forest habitat will be beneficial for biodiversity conservation.

### EFFECTS OF RIPARIAN RESERVE STRUCTURE AND LANDSCAPE CONTEXT

Our results reinforce the importance of enforcing a protected riparian zone where it is already present in legislation and introducing similar legislation in regions where it is absent. However, our data suggest that increasing the protected area to 50 – 80 m on each side of the river is necessary to retain levels of species richness and diversity found in logged forest. For Sabah, these recommendations are wider than the current legal requirement of 20 m and may only be a feasible reality for wider rivers. Therefore, we suggest that multiple tiered recommendations for rivers of different sizes (Darveau *et al.* 2001; Hannon *et al.* 2002; Barclay *et al.* 2017) will be most appropriate, so that wider reserves are maintained in the most critical locations. Recommendations of 50 to 80 are still within the range recommended by the RSPO and the legal requirements of other major oil palm production nations such as Indonesia and Ghana. In addition, specifications for a minimal basal area or percentage canopy cover may also be beneficial, as we found that the vegetation complexity of the reserves can increase the functional diversity of dung beetles and may therefore also affect the important ecological functions they support.

Our results also indicate that landscape context influences the conservation value of riparian reserves. The positive relationship between dung beetle (alpha) diversity and the amount of forest in the area surrounding riparian reserves is consistent with evidence from the neotropics that the surrounding matrix impacts dung beetle communities in forest fragments (Barnes *et al.* 2014). Retaining connections between riparian reserves and forest left on steep slopes could provide synergistic benefits to the communities of both types of forest fragment. community composition changed with distance from large areas of logged forest, as has been found for both birds and dung beetles in riparian forests within neotropical plantations (Hawes *et al.* 2008; Barlow *et al.* 2010). However, we found that many forest dependent species were still present >14 km from a large forested area. Moreover, even though abundance, biomass, and functional diversity were lower in the riparian reserves than in the logged forest, distance from logged forest did not have further negative impacts on these aspects of the dung beetle community.

We were not able to collect data on the extent to which a direct connection to a large area of forest affects the conservation value of riparian reserves, as this landscape is at the edge of the expanding area of oil palm and all riparian corridors were still connected to the logged forest. However, connectivity to larger areas of forest has been shown to impact the communities within riparian reserves in the neotropics (Lees & Peres 2008) and may well have a similar effect in Southeast Asia. Studying riparian reserves in a more homogenous landscape (e.g. at lower elevations where the flatter terrain means that there are fewer forest fragments remaining), where corridors do not directly connect to larger forest areas, would allow us to further study the effects of forest connectivity on the conservation value of riparian reserves.

As our study area is part of a relatively newly converted landscape on the frontier of conversion to oil palm it is possible that the riparian reserve communities are still changing in response to the initial land conversion. We were not able to obtain precise measures of the time since the reserves were isolated as parts of the landscape were previously timber plantations under different management. However, data from satellite images indicates that the majority of deforestation occurred in the 15 years preceding our study (Hansen *et al.* 2013). As it can take > 25 years for extinction debts to be paid (Stouffer *et al.* 2011; Gibson *et al.* 2013) the riparian reserves may therefore still be declining in richness. Obtaining data on how these communities change over the next decade will help establish whether these sites are representative of the long term conservation value of riparian reserves.

These considerations of landscape design and riparian zone management are of clear importance for countries where oil palm plantations are expanding, but also of increasing importance as much of the oil palm industry in Southeast Asia is reaching a replanting phase (Snaddon, Willis & Macdonald 2013). In addition to protecting existing reserves during replanting, there is a great opportunity to put in place restoration plans to rehabilitate riparian zones that were not previously sufficiently protected.

It is also important to emphasise that even though the species richness in the reserves can achieve levels comparable to that in logged forest, the beetle abundance still remains much lower than in logged forest. Whilst we can provide recommendations for maximising the biodiversity within riparian reserves, this strategy is valuable only where plantations are a necessity, and is not comparable to the conservation of larger areas of continuous forest (Edwards *et al.* 2010, 2011; Slade, Mann & Lewis 2011).

### RIPARIAN RESERVES AS CORRIDORS

Our FIT data suggest that there is no net flow of beetles from forest into the reserves. The lack of difference between beetles sampled on different sides of the FITs could be because there are equal numbers of beetles moving in and out of the reserves. However, this result may also merely indicate that beetles in riparian forest have complex flight paths, resulting in equal captures on both sides of the FITS and masking an underlying net flow of individuals out of the forest. We are not aware of this method being widely used to test for movement of invertebrates through linear strips of forest, although our results are non-significant, we feel that it is valuable to present them here to contribute to the future development of techniques to assess the movement and spatial habitat use of dung beetles in tropical forests.

### CONCLUSIONS

Riparian buffers retain many of the species found in large areas of native forest, and should be enforced where relevant legislation exists. Similar legislation should be introduced in oil palm growing regions of south east asia where it is not currently present, and wherever possible within Sabah, we recommend reserves of 50 - 80 m should be protected. Minimising distances between habitat fragments during conversion to oil palm, and maximising the number of trees and canopy cover within remaining fragments should also help retain biodiversity in oil palm landscapes.

## Acknowledgements

We are grateful to EPU Malaysia and Sabah Biodiversity council for research permissions. This work was carried out under the EPU permit UPE 40/200/19/2711 and Sabah Biodiversity Council access license JKM/MBS.1 000-2 12(38). Benta Wawasan Sdn Bhd provided access to field sites and palm yield data. The SAFE project coordinators (Ed Turner, Johnny Larenus and MinSheng Khoo) and several SAFE project research assistants provided logistical support for data collection. cLG was supported by a NERC DTG studentship.

